# Identifiability of speciation times under the multispecies coalescent

**DOI:** 10.1101/2020.11.24.396424

**Authors:** Laura Kubatko, Alexander Leonard, Julia Chifman

**Author notes:** Corresponding author (Laura Kubatko). Email addresses:* (Alexander Leonard), (Julia Chifman).

## Abstract

The advent of rapid and inexpensive sequencing technologies has necessitated the development of computationally efficient methods for analyzing sequence data for many genes simultaneously in a phylogenetic framework. The coalescent process is the most commonly used model for linking the underlying genealogies of individual genes with the global species-level phylogeny, but inference under the coalescent model is computationally daunting in the typical inference frameworks (e.g., the likelihood and Bayesian frameworks) due to the dimensionality of the space of both gene trees and species trees. Here we consider estimation of the branch lengths in fixed species trees with three or four taxa, and show that these branch lengths are identifiable. We also show that for three and four taxa simple estimators for the branch lengths can be derived based on observed site pattern frequencies. Properties of these estimators, such as their asymptotic variances and large-sample distributions, are examined, and performance of the estimators is assessed using simulation. Finally, we use these estimators to develop a hypothesis test that can be used to delimit species under the coalescent model for three or four putative taxa.

## 1. Introduction

The multispecies coalescent (MSC) model is used to describe the process by which nucleotide sequence data arise from a species tree. The model incorporates two distinct processes: the coalescent process [1, 2, 3] is used to generate gene trees given the species tree, and standard nucleotide substitution models [4] are used to generate sequence data along the gene trees for each gene. Over the past 20 years, many methods for estimation of species trees under the MSC have been proposed (see, e.g., Liu et al. (2009) or Kubatko (2019) for reviews of these methods). One reason that so many different methods have been proposed is that standard approaches to estimation, such as those based in the likelihood or Bayesian frameworks, are computationally intensive and often infeasible for data sets of realistic size. Study of properties of the MSC, such as identifiability of parameters, is thus particularly important for developing effective methods for inference.

Figure 1 shows two four-taxon species trees for which the leaves of the trees, labeled *a, b, c*, and *d*, represent present-day species for which DNA sequence data from many loci are observed. The shaded internal branches of the tree, labeled *P*_1_, *P*_2_, and *P*_3_, represent hypothetical common ancestral populations for the species at the leaves. The parameters *τ*_1_, *τ*_2_, and *τ*_3_ are the times from the present back to various speciation events, events in which an ancestral population gives rise to two distinct descendants, indicated by dotted lines. From an inference standpoint, we wish to estimate both the tree topology (i.e., the branching pattern) and the speciation times *τ*_*i*_, *i* = 1, 2, 3. Figure 1 also depicts gene trees as red lines embedded within a given species tree. These trees represent the phylogenetic histories, which we will denote by *h*, of the lineages sampled from species *a, b, c*, and *d* for a particular gene. The parameters *t*_1_, *t*_2_, and *t*_3_ are the times of coalescent events, events in which two lineages first share a common ancestor, measured from the most immediately descendant speciation event. For example, in Figure 1 (left panel) the second coalescent event occurs in population *P*_3_ and it represents the time, denoted by *t*_2_, from the speciation event that resulted in two distinct populations *P*_1_ and *P*_2_ backward until the lineages *c* and *d* share a common ancestor. Note that the coalescent event described in the previous sentence could instead have happened in population *P*_1_, resulting in the same gene tree topology but different coalescent times. Thus, to make this distinction clear, we will associate with each gene tree history *h* a vector of coalescent times **t**_*h*_ corresponding to specific intervals between speciation events. We denote this by a pair (*h*, **t**_*h*_). We begin by defining the probability model for general species phylogenies with any number of tips, before returning to the four-taxon case to establish our main results about species tree branch lengths.

**Figure 1.**
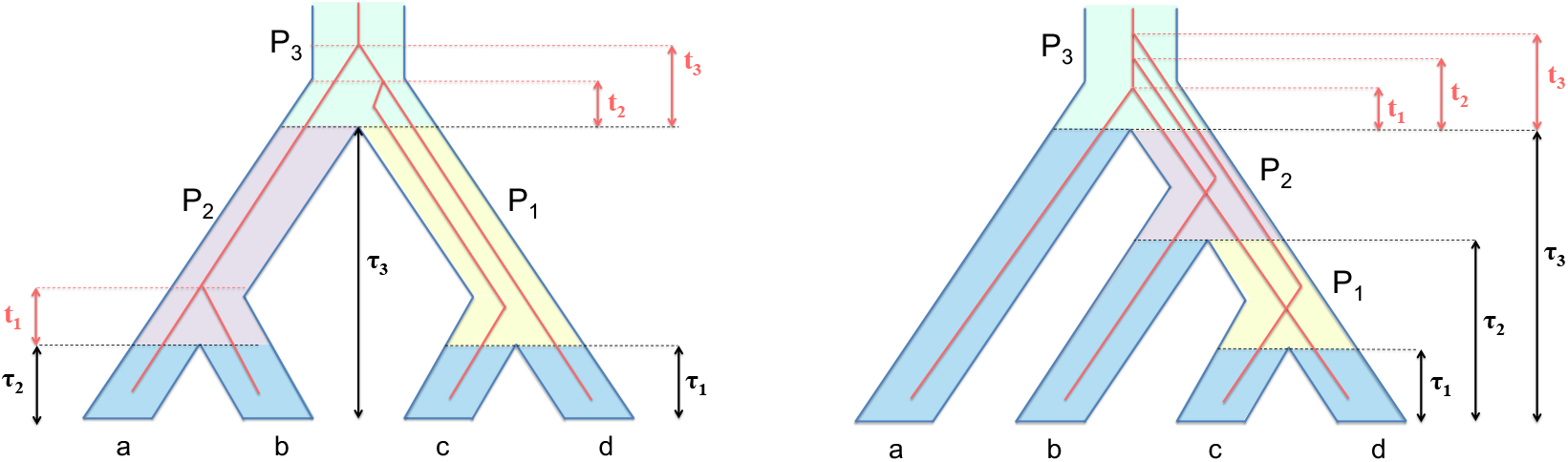
Model symmetric (left) and asymmetric (right) four-taxon species trees with speciation times denoted by *τ*_1_, *τ*_2_, and *τ*_3_. Each speciation time is measured from the present time (the tips of the tree) backward. Note that the definitions of *τ*_1_ and *τ*_3_ are the same in both panels (i.e., *τ*_1_ is the speciation events that separates species c from species d, and *τ*_3_ is the initial speciation event), while the definition of *τ*_2_ differs between the two trees due to their differing shapes. Nested within each of the species trees is an example gene tree history (red lines) that shows the sequence of coalescent events among the sampled lineages for the gene represented. The times of the coalescent events are measured from the immediately descendent speciation event and are denoted by *t*_1_,*t*_2_, and *t*_3_.

Let *M* be the number of species under consideration, and label the species 1, 2, …, *M*. Suppose that *m*_*j*_ individuals are sampled for species *j* so that the total number of individuals being analyzed is 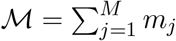 Denote the conditional probability density of gene tree history *h* and associated vector of coalescent times **t**_*h*_ by *f*_(*h*,**t**_*h*_)|(*S,τ*)_, where the conditioning is on species tree topology *S* and vector of speciation times *τ* (a description of how this density is computed can be found in [7]). Define a site pattern as an assignment of states *i*_1_*i*_2_ · · · i_ℳ_ to the ℳ tips of the tree so that for each *k* = 1, 2, …, ℳ, *i*_*k*_ ∈ {*A, C, G, T*}. The probability of site pattern *i*_1_*i*_2_ · · · *i*_ℳ_ arising from *gene tree history* (*h*, **t**_*h*_) is denoted by by 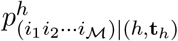. We note that this probability is the typical phylogenetic likelihood for a gene tree, which is computed according to one of the standard nucleotide substitution models. The probability that site pattern *i*_1_*i*_2_ *· · · i*_ℳ_ arises from the *species tree* is then given by

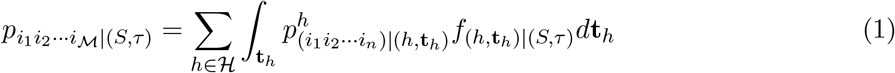

where the sum is over all ℋ of the possible gene tree histories with the corresponding branch lengths **t**_*h*_ integrated out. Full details of the calculations are given in [8].

Equation (1) assumes that each site in the sequence alignment is an independent and identically distributed observation from the MSC model. In other words, each site is a draw from the distribution of gene trees given the species tree under the MSC, with mutation along the sampled gene tree then occurring following one of the standard nucleotide substitution models. We refer to data of this type as *coalescent independent sites* (CIS), and contrast this data type with SNP data, which do not include constant sites. With the model specified here, a data set consisting of CIS can be seen as a sample from the multinomial distribution, with 4^ℳ^ possible categories (i.e., the number of possible sites patterns) and with category probabilities given by the site pattern probabilities.

We note that for a fixed species tree, the site pattern probabilities are functions of the species tree branch lengths (the *τ* s in the equation above), as well as the effective population sizes. In the empirical setting in which data are observed and the goal is to estimate the tree topology, branch lengths, and effective population sizes, a necessary first step is to establish that these model parameters are identifiable. Identifiability of the species tree topology (but not the branch lengths) was proved by Chifman and Kubatko (2015) for species trees that satisfy the molecular clock for the GTR+I+Γ substitution model and all submodels thereof. Long and Kubatko (2019) proved identifiability for non-clock species tree topologies and for tree topologies in which the effective population sizes vary throughout the tree. However, identifiability of other model parameters, such as the branch lengths, has not been formally established in any of these cases.

Here, we show that the branch lengths in three- and four-taxon species trees are identifiable under the multispecies coalescent with the JC69 model [10] for the setting considered by [8] in the case in which one lineage is sampled per species and a single effective population size is assumed for the whole tree. The method used to establish indentifiability leads to a natural estimator of the branch lengths for the case of four taxa that can be computed analytically. Variance estimates are also available analytically in this case. In Section 2, we provide a proof of the identifiability of the branch lengths. While our proof uses the site pattern probabilities computed in Chifman and Kubatko (2015), this result was not provided in our previous work. In Section 3, we derive novel formulas for estimators of the branch lengths that are motivated by the proof presented here, along with formulas for the variances of these estimators. Corresponding results for the case of three taxa are given in Appendix C. Section 4 uses simulation to evaluate the performance of these estimators for simulated data. In Section 5, we describe an application of the branch length estimators to the problem of species delimitation, and we apply this method to an empirical data set in Section 6. We conclude in Section 7 with a discussion of the issues that arise when applying these estimators to larger trees, as well as some applications in which these estimators can be used.

## 2. Identifiability of the species tree branch lengths

We begin by considering the case in which *M* = 4 and *m*_*j*_ = 1 for *j* = 1, 2, 3, 4, i.e., a four-taxon species tree with one lineage sampled per species. In this case, there are 15 possible site patterns given by

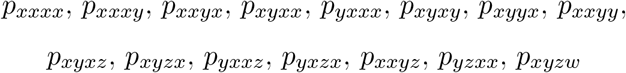

where *x, y, w*, and *z* denote distinct nucleotides. Let **Y** = (*Y*_1_, *Y*_2_, …, *Y*_15_) denote the vector of counts of these site patterns in an observed data set, and let **p** = (*p*_1_, *p*_2_, …, *p*_15_) denote their probabilities. Then for a sample of *n* CIS under the assumptions of the MSC, **Y** *∼* Multinominal(*n*, **p**), and thus we have the following:

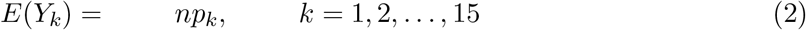

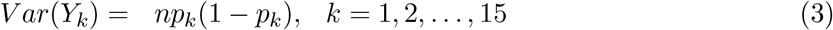

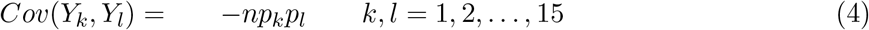

Under the JC69 model, many of these site patterns will have equal probabilities. For example, Chifman and Kubatko (2015) show that for the rooted symmetric four-taxon species tree, these 15 site pattern probabilities can be further reduced to 9 distinct site pattern probabilities, while for the rooted asymmetric four-taxon species tree, there are 11 distinct site pattern probabilities. Moreover, they give formulas for these site pattern probabilities for both the symmetric and asymmetric four-taxon species trees as a function of the species tree branch lengths (*τ*) under the assumption of a single effective population size, *θ* for the entire tree. As is standard in species tree modeling using the coalescent process, we define the effective population size parameter *θ* = 4*N*_*e*_*µ*^*′*^, where *N*_*e*_ is the population size and *µ*^*′*^ is mutation rate in units of substitutions per site per generation in the DNA sequence. The branch lengths in the species tree are given in coalescent units, which are the number of 2*N*_*e*_ generations. These site pattern probabilities can be used to establish identifiability, as shown below.

*Case 1: The Symmetric Species Tree*. Consider first the case of the symmetric species tree (Figure 1). The 15 site pattern probabilities listed above can be reduced to 9 distinct site pattern probabilities in this case:

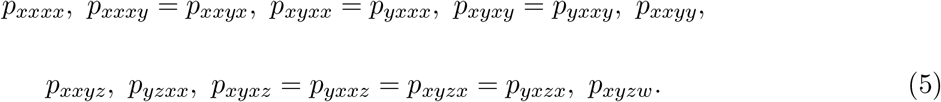

Chifman and Kubatko (2015) showed that for *i*_*k*_ ∈ {*A, C, G, T*}, *k* = 1, 2, 3, 4, these site pattern probabilities can be written as

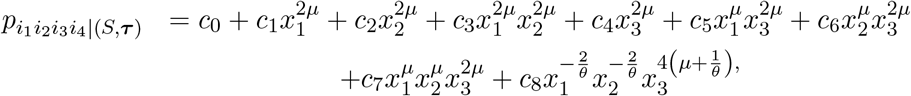

where 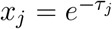 for *j* = 1, 2, 3, *µ* = 4*/*3 for the JC69 model (so that gene tree branch lengths are measured as the expected number of mutations per site), and the coefficients *c*_*i*_, *i* = 0, 1, 2, …, 8 are given in Appendix A.

Let (**C**_**s**_)_9×9_ be the matrix of coefficients given in Appendix A, and note that the above expressions for the site pattern probabilities can be written in matrix form as

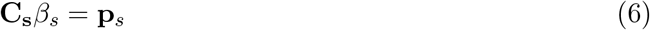

where

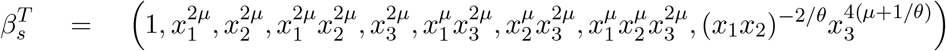

and

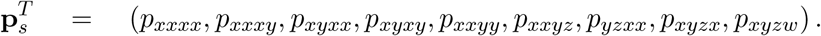

Note that **C**_**s**_ is full rank, and thus the speciation times *τ*_1_, *τ*_2_, and *τ*_3_ can be identified. In particular, we solve Equation (6) for *β*_*s*_. Letting 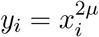, for *i* ∈ {1, 2, 3}, the unique solution is

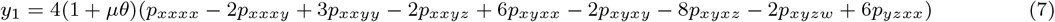

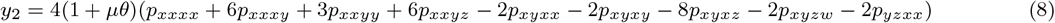

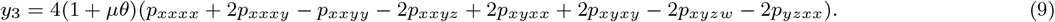

Recalling that 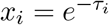, we convert back to speciation times using:

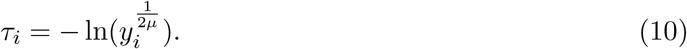

*Case 2: The Asymmetric Species Tree*. We now consider the case of the asymmetric species tree (Figure 1). In this case, the 15 possible site pattern probabilities can be reduced to 11 distinct probabilities:

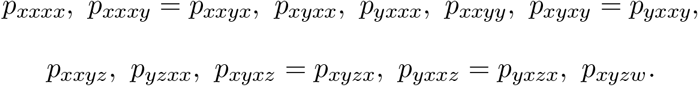

Chifman and Kubatko (2015) showed that for *i*_*k*_ ∈ {*A, C, G, T*}, *k* = 1, 2, 3, 4, the site pattern probabilities can be written as

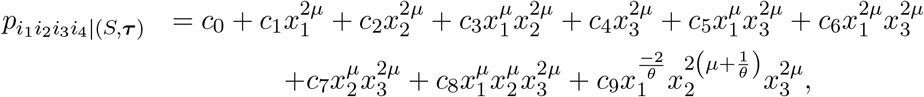

where 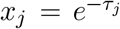 for *j* = 1, 2, 3, *µ* = 4*/*3 for the JC69 model, and the coefficients in the above expression are given in Appendix A. Letting (**C**_**a**_)_11×10_ be the matrix of coefficients, the above expressions for the site pattern probabilities can be written as

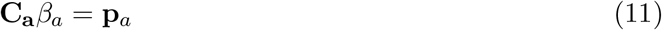

where

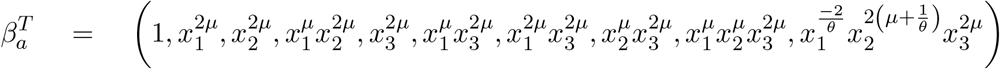

and

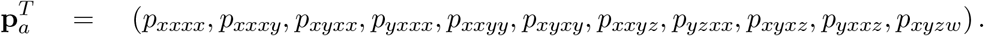

Since matrix **C**_**a**_ has full column rank, we can solve Equation (11) for β_*a*_ using the left-inverse of **C**_**a**_ to obtain the following estimators

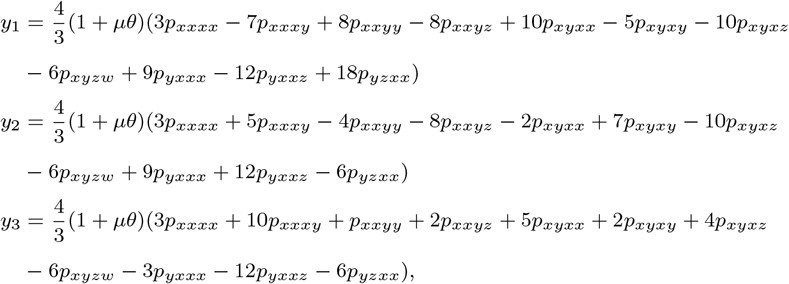

where 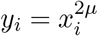 for *i ∈ {*1, 2, 3*}*. The following expression for the left inverse was used (**C**_**a**_^*T*^ **C**_**a**_)^−1^**C**_**a**_^*T*^. We solve for *τ*_*i*_ as before:

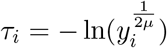

*Identifiability of the speciation times*. The work above establishes that the species tree branch lengths are identifiable for both the symmetric and asymmetric species trees under the MSC for the JC69 model for a four-taxon species tree. We note that we used the JC69 model for our initial verification of identifiability because closed-form expressions for the site patterns probabilities are available. But we note that for any sub-model of GTR+I+Γ expressions analogous to Equations (6) and (11) can be written once the parameters of the substitution model are specified. However, it is not immediately obvious under what models and conditions the system **p** = **C***β* will be consistent, and this will require careful investigation. If the system is consistent, then it is possible that for some sub-models the matrix **C** will have full column rank, in which case the left inverse exists and a unique solution can be computed. It is also possible that the system **p** = **C***β* will be underdetermined for some sub-models but still consistent. In this case there are infinitely many solutions and the pseudoinverse can be used to find the minimum length solution of the system. If there are parameter choices for which the system is inconsistent, the pseudoinverse can be used again and, in this case, provides the least-squares solution of minimum length.

### 3. Moment-based estimators of the branch lengths

The previous section provides explicit analytic formulas for the branch lengths for both the symmetric and asymmetric rooted four-taxon species trees under the MSC for the JC69 model. In this section, we show how the variances of these estimators can be estimated.

To illustrate the main ideas, consider the case of the symmetric species tree in Figure 1 and suppose that our interest is in estimating 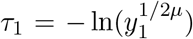. Let **Y**_**s**_ = (*Y*_*s*,1_, *Y*_*s*,2_, …, *Y*_*s*,9_) be the observed counts of the site patterns corresponding to each of the patterns listed in (5) above, and recall that under the MSC model for a sample of *n* CIS, **Y** ∼ Multinomial(*n*, **p**_*s*_). Thus, for large *n*, we have

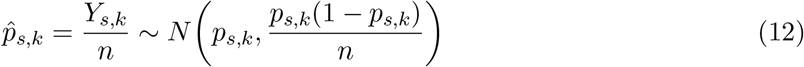

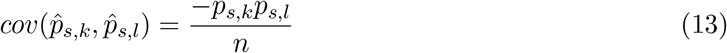

for *k, l* = 1, 2, …, 9.

To estimate 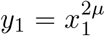, we can substitute the empirical estimates 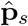 into Equation (7) for **p**_*s*_. We denote this estimator *ŷ*_1_. We can also find an expression for the asymptotic variance of this estimate by taking the variance in Equation (7) and substituting the appropriate variance and covariance terms from (12) and (13) to get

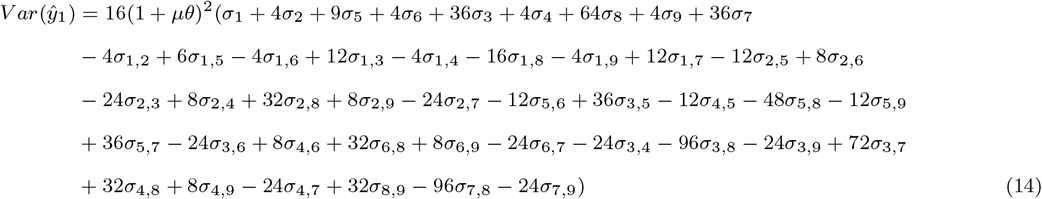

where 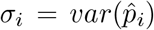 and 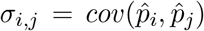 We estimate all of the quantities in the expression above using the empirical frequencies to obtain 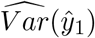, which is an estimator of this asymptotic variance.

Similar expressions can be derived for all other branch lengths in the both the symmetric and the asymmetric species tree. We provide these expressions in Appendix B. Note that it is easy to convert *ŷ*_*i*_ to the original scale of *τ*, using

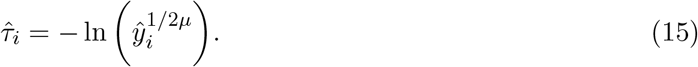

When *n* is large, *ŷ*_*i*_ will be approximately normally distributed with mean *y*_*i*_ and variance that can be approximated by 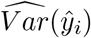. Furthermore, the asymptotic normal distribution can be used to construct a (1 *− α*) × 100% confidence interval using

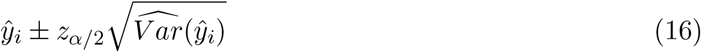

where *z*_*α/*2_ is the value such that Φ(*z*_*α/*2_) = 1 *− α/*2 where Φ is the cumulative distribution function of the standard normal distribution. This interval can be transformed to the original scale of *τ* by applying Equation (15) to both endpoints.

To summarize, the estimators derived here can be used to obtain unbiased point estimates of the speciation times, as well as corresponding large-sample confidence intervals. It is to be noted that all of this can be achieved with closed-form equations, i.e., no optimization is required, and thus these quantities can be computed instantaneously. In the next section, we use simulation to show that these estimators behave as expected.

## 4. Simulations

We examine the performance of the estimators using simulation. The overall simulation design is as follows:

1. Simulate *n* CIS from Multinomial(n,**p**_**s**_) or Multinomial(n,**p**_**a**_).
2. Compute *ŷ*_*i*_ and 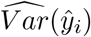 for *i* = 1, 2, 3.
3. Construct 95% confidence intervals for *τ*_*i*_, *i* = 1, 2, 3 by transforming the endpoints of the interval in Equation (16) using (15).
4. Repeat steps 1-3 *B* times.

In Step 1, we must specify values for all of the parameters. We consider the settings below for both the symmetric and the asymmetric model trees in Figure 1. The branch lengths *τ*_*i*_, *i* = 1, 2, 3 are given in coalescent units (number of 2*N*_*e*_ generations) and *θ* is the effective population size.

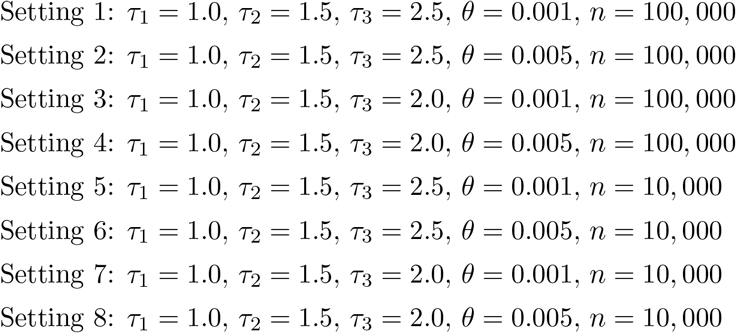

Settings 1-4 consider a total of *n* = 100, 000 CIS, which is a moderately large dataset size that is typical of many genome-scale phylogenomics studies. Settings 5-8 consider a smaller dataset size, with only *n* = 10, 000 CIS. In odd numbered settings, the effective population size parameter *θ* is set to 0.001, while in the even numbered settings, *θ* = 0.005. These represent typical values that are on the lower (odd settings) and upper (even settings) range of values commonly observed. The values of *τ*_1_ and *τ*_2_ are held constant in all of the settings at values that are typical but challenging, since they represent relatively short branch lengths. The value of *τ*_3_ is varied to provide both a very difficult problem (*τ*_3_ = 2.0, creating an internal branch of length 0.5 coalescent units; settings 3, 4, 7, and 8) or a slightly easier problem (*τ* = 2.5, creating a branch of length 1.0 coalescent units; settings 1, 2, 5, and 6). For all simulation settings, *B* = 25, 000.

For each simulation setting, histograms of the estimates *ŷ*_*i*_ and 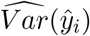 are constructed (Figures 2 and 3 show these histograms for Setting 1; figures for all other settings are shown in the Supplemental Material). The thick gray line corresponds to the value of the parameter estimated across the *B* simulation replicates (i.e., for the histogram of the estimates *ŷ*_*i*_, the gray line corresponds to the mean of the estimates across the *B* replicates; for the histogram of the estimates 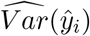, the gray line corresponds to the sample variances of the estimates *ŷ*_*i*_ across the *B* replicates). The blue lines in each plot give the “true” values of the parameters, i.e., the values computed by plugging the true site pattern probabilities into the expressions for the estimators. We note that, in general, it is easier to estimate a mean than a variance and thus the blue lines and gray lines agree more closely for the plots of *ŷ*_*i*_. Additionally, we plot 95% confidence intervals for *τ*_*i*_, *i* = 1, 2, 3, for the first 100 replicates, with a red line corresponding to the true value of *τ*_*i*_. The observed portion of the *B* replicates that contain the true value is given in the plot. We show results for Setting 1 here for both the symmetric (Figure 2) and asymmetric (Figure 3) species trees. The Supplemental Material contains the plots for the other simulation settings. In addition, an R Markdown file that will allow the reader to reproduce these simulation results as well as examine different settings is available at https://github.com/lkubatko/MOMSpeciationTimes.

**Figure 2.**
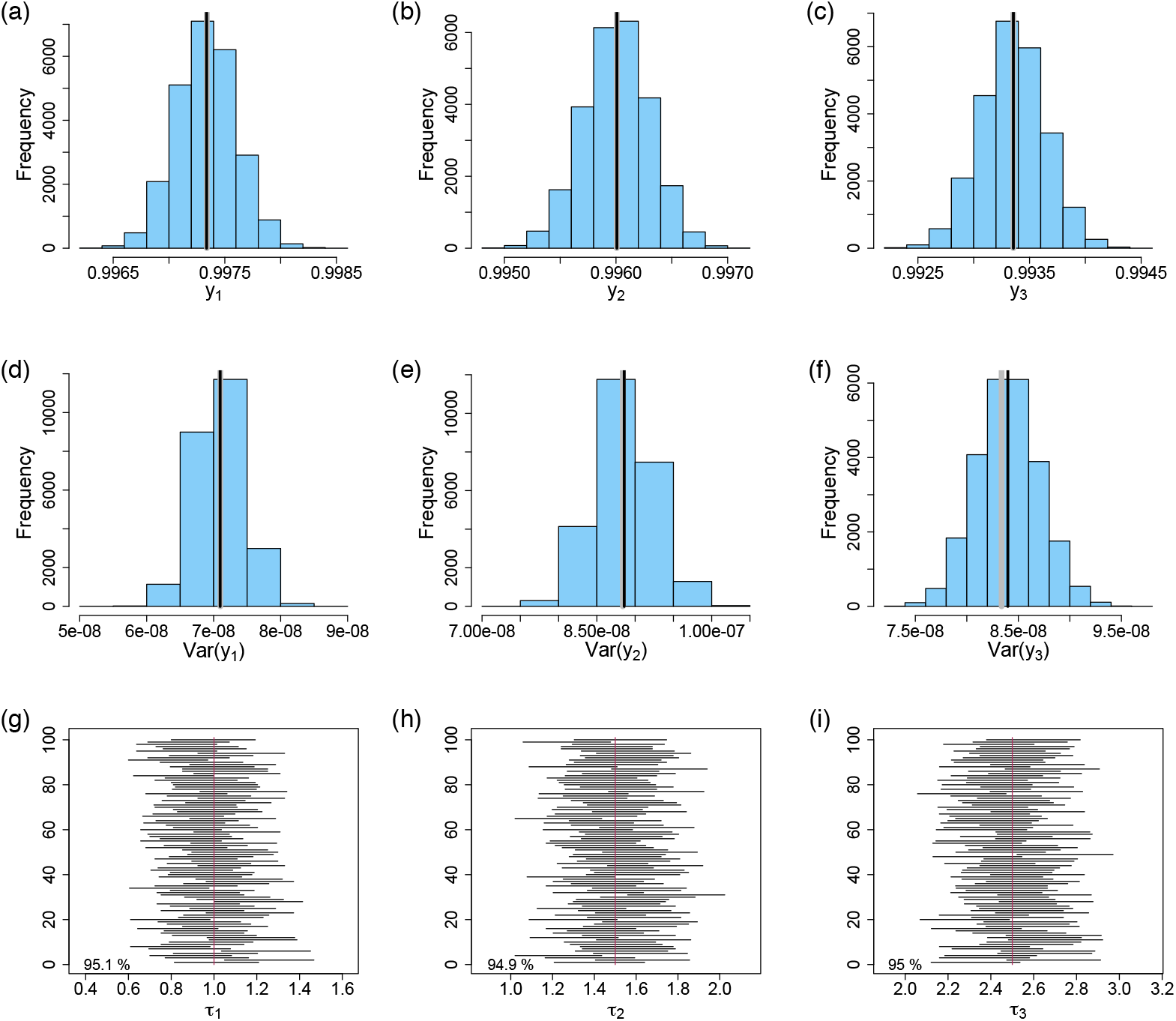
Results of the simulation for Setting 1 for the symmetric species tree in Figure 1. The first row shows histograms of (a) 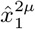, (b)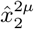, and (c)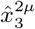. The second row shows histograms of (d)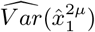, (e)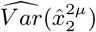, and (f)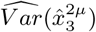. In the first two rows, the gray line corresponds to the value obtained by summarizing the simulated values, and the blue lines corresponds to the true values (see text for details). The third row plots the first 100 of the simulated 95% confidence intervals for the (g) *τ*_1_, (h) *τ*_2_, and (i) *τ*_3_. The red line indicates the true value, and the number in the bottom left corner is the portion of all simulation replicates for which the confidence interval includes the true value.

**Figure 3.**
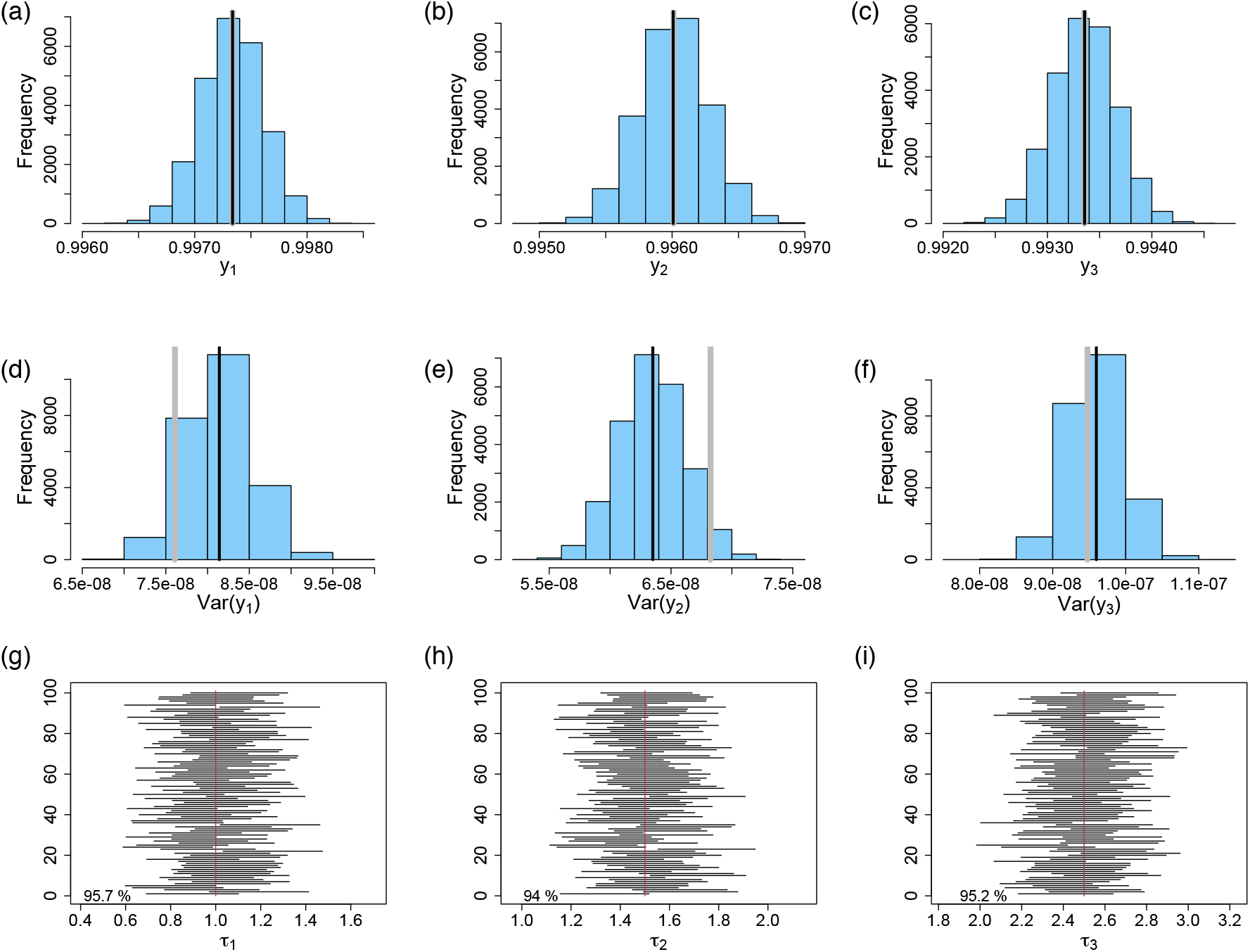
Results of the simulation for Setting 1 for the asymmetric species tree in Figure 1. The first row shows histograms of (a)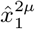, (b)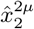, and (c)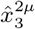. The second row shows histograms of (d)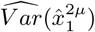, (e)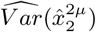, and (f)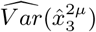. In the first two rows, the gray line corresponds to the value obtained by summarizing the simulated values, and the blue lines corresponds to the true values (see text for details). The third row plots the first 100 of the simulated 95% confidence intervals for the (g) *τ*_1_, (h) *τ*_2_, and (i) *τ*_3_. The red line indicates the true value, and the number in the bottom left corner is the portion of all simulation replicates for which the confidence interval includes the true value.

An examination of subfigures (a) - (c) in Figures 2 and 3 indicates that the sampling distribution for *ŷ*_*i*_, *i* = 1, 2, 3, appears to be normally distributed, as expected, and that *y*_*i*_ is unbiasedly estimated. Subfigures (d) - (f) in each panel show the distribution of estimated variances, again indicating that the variance is accurately estimated. We note that the value estimated from the simulated values of *ŷ*_*i*_, represented by the gray line in the figures, is further from both the simulated estimates of the variance and the true value in the case of asymmetric trees. This is because a larger number of simulation replicates is needed to accurately estimate variances than means and because the variances are quite small in this setting. Because the variance for the asymmetric tree contains more terms than does the variance for the symmetric tree (see the expressions in Appendix B), the variance is more difficult to estimate by using the variance of the estimates in this case. However, the accuracy of our estimated variances is supported by the strong performance of our confidence intervals for the speciation times. In particular, we note that all 95% confidence intervals show coverage probabilities very close to 95% – across all 16 simulation settings and all three parameters, the empirical coverage was always between 93.7% and 96.0%, suggesting that our estimates of both the branch lengths and their variances are very accurate.

## 5. Application to species delimitation

The estimators derived in the previous section can be applied to the problem of species delimitation, in which one seeks to determine whether two sets of taxa are members of distinct species (vs. members of the same species). For example, consider the symmetric species tree in Figure 1, and suppose that we are interested in addressing the question of whether lineages sampled from tip *c* form a distinct species from those sampled from tip *d*. One way to ask this question is to ask whether the speciation time *τ*_1_ is significantly larger than 0, since if *c* and *d* are not distinct species, then it should be the case that *τ*_1_ = 0.

Formally, we propose to test the hypotheses

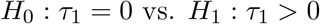

using the test statistic

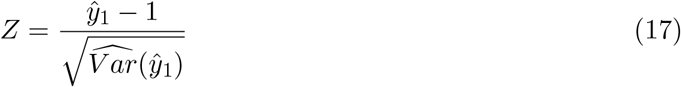

where we subtract one in the numerator because *τ*_1_ = 0 implies *y*_1_ = 1. When the number of sites, *n*, is large, *Z* ∼ *N* (0, 1) when *H*_0_ is true, and we reject *H*_0_ when *Z > z*_*α*_.

We test the performance of this method using simulation. For Settings 1, 2, 5, and 6, we generated *B* = 1, 000 data sets for *τ*_1_ *∈ {*0.025, 0.05, 0.075, 0.1, 0.125, 0.15, 0.175, 0.2, 0.3, 0.4, 0.5, 0.6, 0.7, 0.8, 0.9, 1.0*}*, and computed the test statistic in expression (17) above. For each data set, we computed the power of the test, the proportion of times that the test rejected *H*_0_ at level *α* = 0.05. The power as a function of *τ*_1_ is plotted in Figure 4. To verify the normal approximation, we also plot the test statistics simulated for three choices of *τ*_1_ (*τ*_1_ = 0, 0.1, 1.0) in Figure S15 in the Supplemental Material.

**Figure 4.**
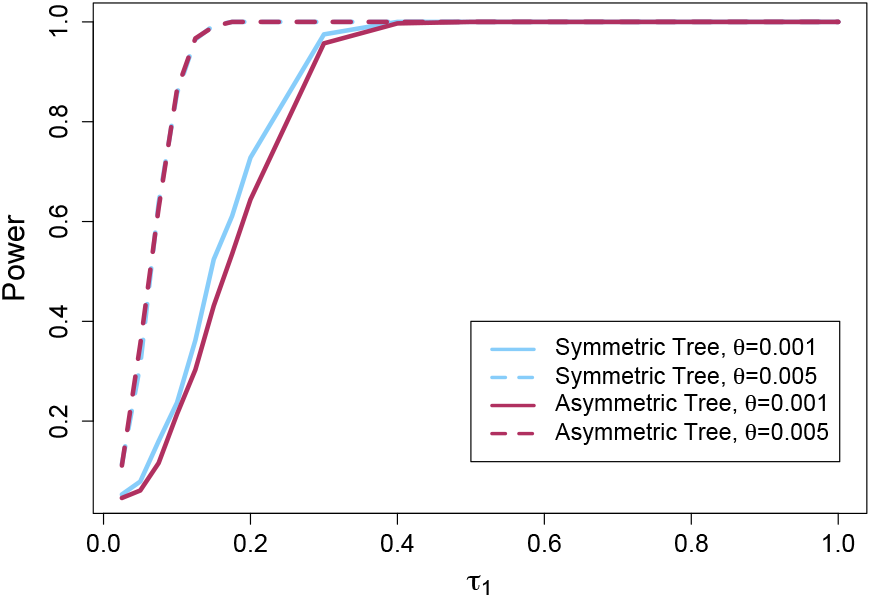
Power of the hypothesis test for both the symmetric (blue) and the asymmetric (red) trees for Setting 1 (*θ* = 0.001, solid lines) and Setting 2 (*θ* = 0.005, dashed lines).

It is easy to see that the test performs well. Figure S15 shows that the test statistics are well-approximated by the normal distribution. The power of the test (Figure 4) increases with increasing *τ*_1_, with the test showing power near 100% for *τ*_1_ *>* 0.1 when *θ* = 0.005 and for *τ*_1_ *>* 0.4 when *θ* = 0.001. The level of the test (the power computed at the null hypothesis value of *τ*_1_ = 0) is also reasonable (0.049 and 0.044 when *θ* = 0.001 for the symmetric and asymmetric trees, respectively, and 0.112 and 0.107 when *θ* = 0.005 for the symmetric and asymmetric trees, respectively), close to the expected value of 0.05 in all cases.

## 6. Empirical Example: *Sistrurus* rattlesnakes

We demonstrate the application of our methodology to an example data set for *Sistrurus* rattlesnakes. Kubatko et al. (2011) analyzed sequence data collected from 19 genes for two species of rattlesnakes, *Sistrurus catenatus* and *S. miliarius*, each of which consists of 3 identified subspecies. Among the questions addressed by these authors was whether one of the subspecies of *S. catenatus* had diverged sufficiently from the other two that it could be considered a distinct species. Kubatko et al. (2011) provided support for this new species designation based on both genetic analyses and morphological distinctiveness. We use the method described in the previous section to assess support for this conclusion.

Referring to the symmetric tree in Figure 1 (left), we consider data for the following subspecies: a = *S. m. miliarius*; b = *S. m. streckeri*; c = *S. c. catenatus*; and d = *S. c. tergeminus*. To test whether our data support *S. c. catenatus* as a distinct species, we test the null hypothesis that *τ*_1_ = 0 using the test statistic in Equation (17). Selecting one individual of each of the species listed above gives the vector of observed site patterns (7909, 127, 46, 0, 103, 9, 1, 0, 0), ordered as in (5). We assume that *θ* = 0.002 and *µ* = 4.0*/*3.0, corresponding to the Jukes-Cantor model. Using expressions (7), (14), (15), and (16), a 95% confidence interval for *τ*_1_ is (2.49, 3.91). The test statistic for the hypothesis test is *z* = −8.94, resulting in a p-value of 1.94 × 10^−19^. Thus, we reject the null hypothesis that *τ*_1_ = 0, meaning that our analysis shows support for *S. c. catenatus* as a distinct species from *S. c. tergeminus* in agreement with the findings of [11].

We also consider estimation of *τ*_2_ and find an estimate of 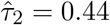 with 95% confidence interval (0.03, 0.85), suggesting a much more recent divergence between subspecies within *S. miliarius*, again consistent with [11]. Our estimates are given here in coalescent units, while [11] reported estimates in millions of years before the present. However, comparing our estimates with theirs (see their Table 5), we find similar results: within *S. catenatus*, a large divergence time is inferred for the separation of subspecies *S. c. catenatus* from *S. c. tergeminus*, while within species *S. miliarius*, the divergence among subspecies is inferred to be much more recent. We note that the analysis in [11] was carried out with a computationally-intensive Bayesian approach requiring hours of computing time, while our analysis can be done with a standard calculator.

## 7. Discussion and conclusions

While our earlier work [8] established identifiability of the species tree topology under the MSC, the work presented here is the first to establish identifiability of the speciation times (or, equivalently, the branch lengths) in a species-level phylogenetic tree under the multispecies coalescent model. In particular, we show that for both the symmetric and asymmetric four-taxon species trees that satisfy the molecular clock, the speciation times (the times of the internal nodes) can be identified for the standard nucleotide substitution models. For the JC69 model [10], we provide specific formulas for estimating these branch lengths, and show that they display the expected large-sample behavior, i.e., they are unbiased and approximately normally distributed. Moreover, simple formulas for estimating the variances of these estimators can be derived. Using properties of the normal distribution, we construct confidence intervals for the speciation times based on these estimators, and use simulation to show that these achieve the expected coverage. We also develop a hypothesis test for speciation that can be used to aid in species delimitation, and we use simulation to show that the test has good power to detect speciation while still achieving the nominal significance level under the null hypothesis.

All of the work here has assumed that the effective population size parameter, *θ*, is known. In fact, the solutions to Equations (6) and (11) provide relationships that this parameter must satisfy and can be used to estimate *θ*, as well, in the case of the JC69 model. However, there is not a closed form solution for *θ* and its solution must be iterated with the values of *τ*. Another assumption in this work is that the species tree satisfies the molecular clock. While this assumption is quite commonly made when the goal is to estimate a species-level phylogeny, it is interesting to consider the case when the clock does not hold. The work of Long and Kubatko (2019) provides site pattern probabilities in the non-clock case, and could be used to examine questions of identifiability in this setting.

Finally, we note that our work here has examined only four-taxon species trees. While one might hope to scale this approach up to larger trees, especially because the estimators derived here consist of formulas that can be rapidly computed, this approach is complicated by the fact that expressions for the site pattern probabilities for trees containing more than four species have not been derived, owing to the computational complexity of this setting (see, e.g., Chifman and Kubatko (2015) or Kubatko (2019) for details). Nonetheless, the site pattern probabilities and multinomial distribution function used here have led to estimation of the speciation times for larger trees using a composite likelihood approach [12]. However, this approach requires optimization of the composite likelihood function, and thus is less computationally efficient than the method presented here. Therefore, the method here is useful for addressing questions about speciation times and species designations when the primary interest is in examining a small collection of closely related taxa.

## 8. Acknowledgements

We thank Elizabeth Allman, John Rhodes, and Seth Sullivant for helpful discussions on this work. L.K. and J.C. were supported by funding from the National Science Foundation (DMS 11-06706).

## Appendix A.

**Table 1:**
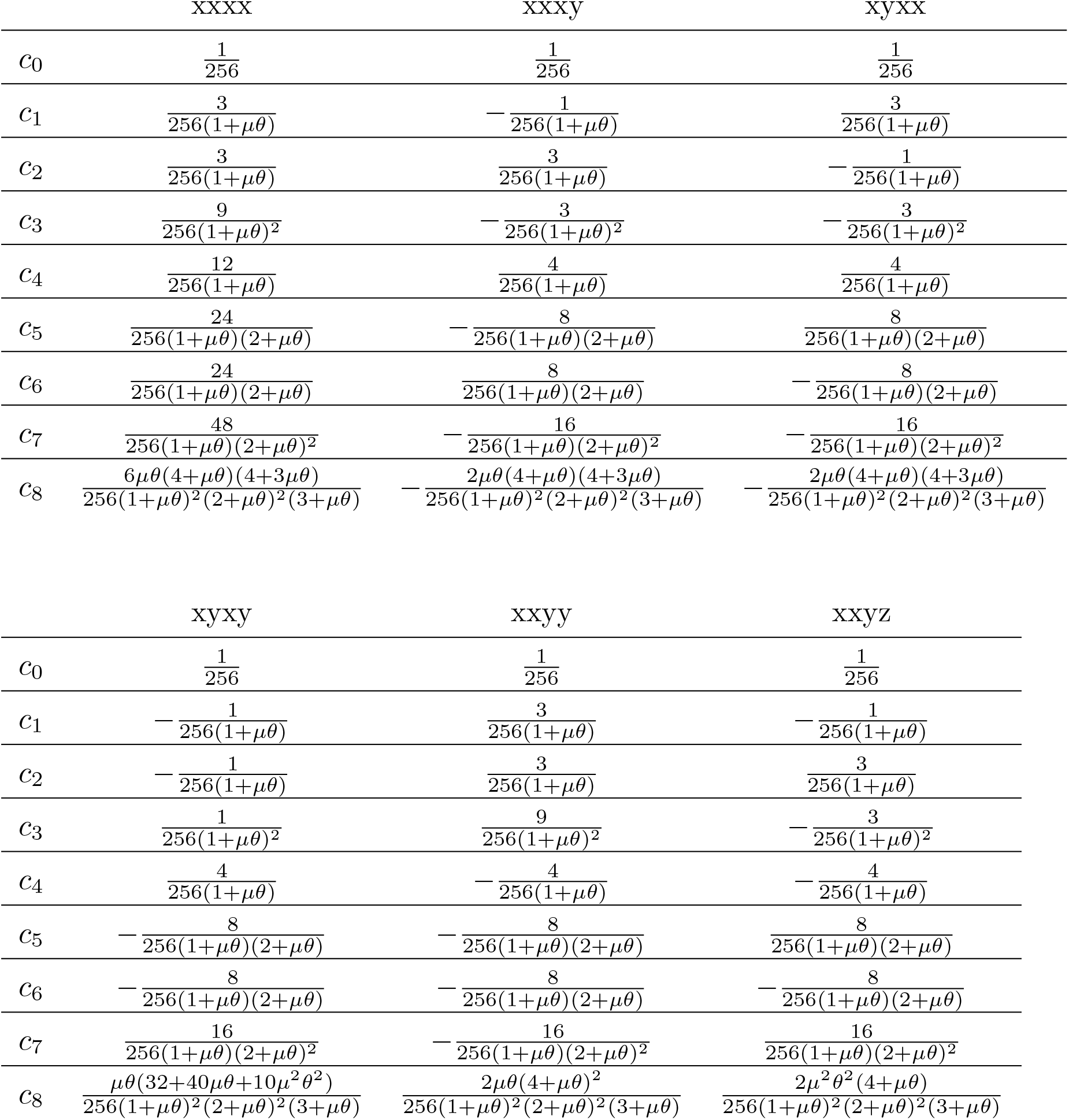
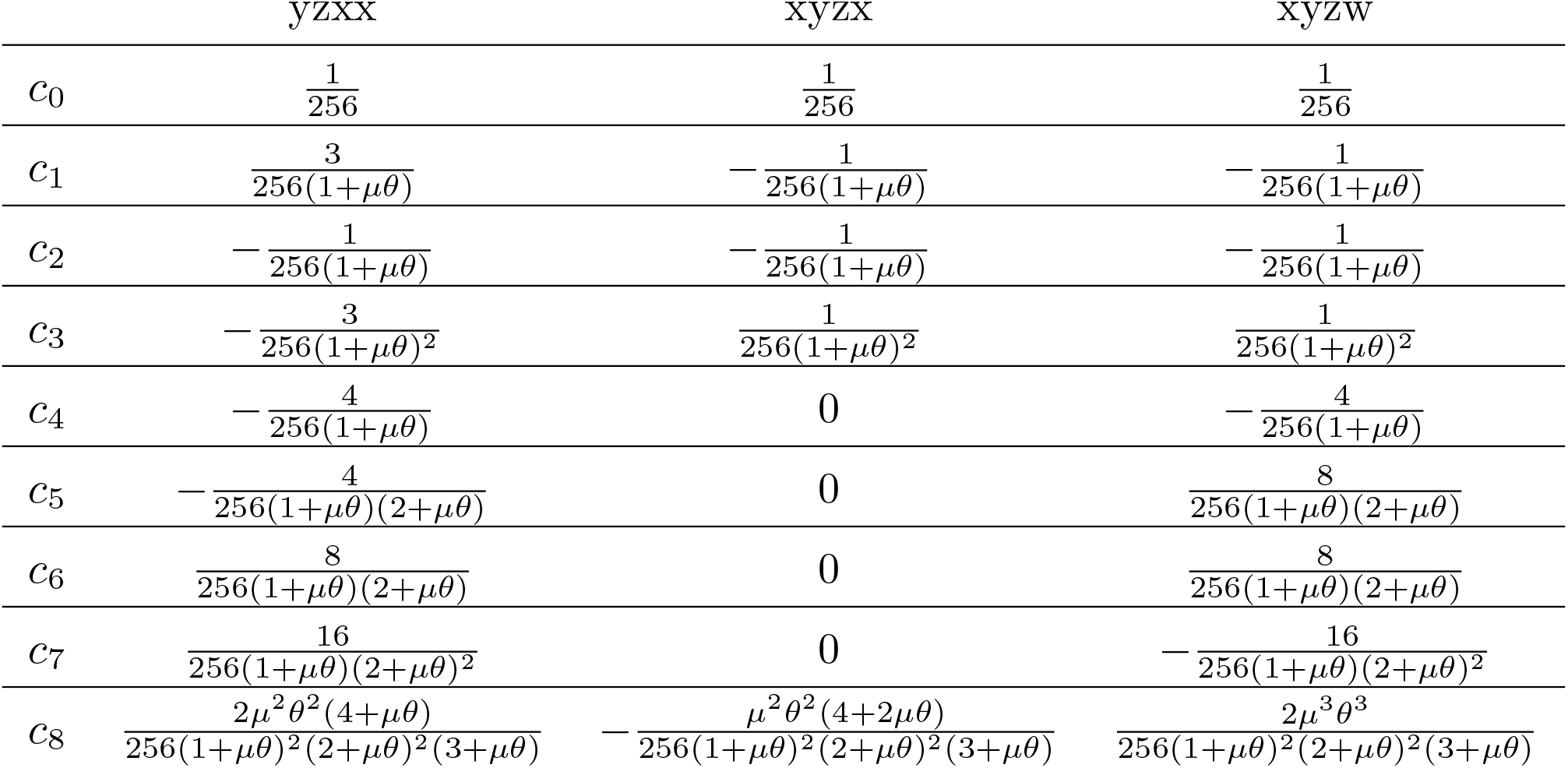
Coefficients for site pattern probabilities: symmetric species 4-leaf tree.

**Table 2:** Coefficients for site pattern probabilities: asymmetric species 4-leaf tree.

|  | xxxx | xxxy | xyxx | yxxx |
| --- | --- | --- | --- | --- |
| $c_0$ | $\frac{1}{256}$ | $\frac{1}{256}$ | $\frac{1}{256}$ | $\frac{1}{256}$ |
| $c_1$ | $\frac{3}{256(1+\mu\theta)}$ | $-\frac{1}{256(1+\mu\theta)}$ | $\frac{3}{256(1+\mu\theta)}$ | $\frac{3}{256(1+\mu\theta)}$ |
| $c_2$ | $\frac{6}{256(1+\mu\theta)}$ | $\frac{2}{256(1+\mu\theta)}$ | $-\frac{2}{256(1+\mu\theta)}$ | $\frac{6}{256(1+\mu\theta)}$ |
| $c_3$ | $\frac{12}{256(1+\mu\theta)(2+\mu\theta)}$ | $-\frac{4}{256(1+\mu\theta)(2+\mu\theta)}$ | $-\frac{4}{256(1+\mu\theta)(2+\mu\theta)}$ | $\frac{12}{256(1+\mu\theta)(2+\mu\theta)}$ |
| $c_4$ | $\frac{9}{256(1+\mu\theta)}$ | $\frac{5}{256(1+\mu\theta)}$ | $\frac{5}{256(1+\mu\theta)}$ | $-\frac{3}{256(1+\mu\theta)}$ |
| $c_5$ | $\frac{6}{256(1+\mu\theta)(2+\mu\theta)}$ | $-\frac{4}{256(1+\mu\theta)(2+\mu\theta)}$ | $\frac{12}{256(1+\mu\theta)(2+\mu\theta)}$ | $-\frac{4}{256(1+\mu\theta)(2+\mu\theta)}$ |
| $c_6$ | $\frac{9}{256(1+\mu\theta)^2}$ | $-\frac{3}{256(1+\mu\theta)^2}$ | $-\frac{3}{256(1+\mu\theta)^2}$ | $-\frac{3}{256(1+\mu\theta)^2}$ |
| $c_7$ | $\frac{24}{256(1+\mu\theta)(2+\mu\theta)}$ | $\frac{8}{256(1+\mu\theta)(2+\mu\theta)}$ | $-\frac{8}{256(1+\mu\theta)(2+\mu\theta)}$ | $-\frac{8}{256(1+\mu\theta)(2+\mu\theta)}$ |
| $c_8$ | $\frac{48}{256(1+\mu\theta)(2+\mu\theta)^2}$ | $-\frac{16}{256(1+\mu\theta)(2+\mu\theta)^2}$ | $-\frac{16}{256(1+\mu\theta)(2+\mu\theta)^2}$ | $-\frac{16}{256(1+\mu\theta)(2+\mu\theta)^2}$ |
| $c_9$ | $\frac{6\mu\theta(4+\mu\theta)(4+3\mu\theta)}{256(1+\mu\theta)^2(2+\mu\theta)^2(3+\mu\theta)}$ | $-\frac{2\mu\theta(4+\mu\theta)(4+3\mu\theta)}{256(1+\mu\theta)^2(2+\mu\theta)^2(3+\mu\theta)}$ | $-\frac{2\mu\theta(4+\mu\theta)(4+3\mu\theta)}{256(1+\mu\theta)^2(2+\mu\theta)^2(3+\mu\theta)}$ | $-\frac{2\mu\theta(4+\mu\theta)(4+3\mu\theta)}{256(1+\mu\theta)^2(2+\mu\theta)^2(3+\mu\theta)}$ |

|  | xxyy | xyxy | xxyz | yzxx |
| --- | --- | --- | --- | --- |
| $c_0$ | $\frac{1}{256}$ | $\frac{1}{256}$ | $\frac{1}{256}$ | $\frac{1}{256}$ |
| $c_1$ | $\frac{3}{256(1+\mu\theta)}$ | $-\frac{1}{256(1+\mu\theta)}$ | $-\frac{1}{256(1+\mu\theta)}$ | $\frac{3}{256(1+\mu\theta)}$ |
| $c_2$ | $-\frac{2}{256(1+\mu\theta)}$ | $\frac{2}{256(1+\mu\theta)}$ | $-\frac{2}{256(1+\mu\theta)}$ | $-\frac{2}{256(1+\mu\theta)}$ |
| $c_3$ | $-\frac{4}{256(1+\mu\theta)(2+\mu\theta)}$ | $-\frac{4}{256(1+\mu\theta)(2+\mu\theta)}$ | $\frac{4}{256(1+\mu\theta)(2+\mu\theta)}$ | $-\frac{4}{256(1+\mu\theta)(2+\mu\theta)}$ |
| $c_4$ | $\frac{1}{256(1+\mu\theta)}$ | $\frac{1}{256(1+\mu\theta)}$ | $\frac{1}{256(1+\mu\theta)}$ | $-\frac{3}{256(1+\mu\theta)}$ |
| $c_5$ | $-\frac{4}{256(1+\mu\theta)(2+\mu\theta)}$ | $-\frac{4}{256(1+\mu\theta)(2+\mu\theta)}$ | $\frac{4}{256(1+\mu\theta)(2+\mu\theta)}$ | $-\frac{4}{256(1+\mu\theta)(2+\mu\theta)}$ |
| $c_6$ | $\frac{9}{256(1+\mu\theta)^2}$ | $\frac{1}{256(1+\mu\theta)^2}$ | $-\frac{1}{256(1+\mu\theta)^2}$ | $-\frac{3}{256(1+\mu\theta)^2}$ |
| $c_7$ | $-\frac{8}{256(1+\mu\theta)(2+\mu\theta)}$ | $-\frac{8}{256(1+\mu\theta)(2+\mu\theta)}$ | $-\frac{8}{256(1+\mu\theta)(2+\mu\theta)}$ | $\frac{8}{256(1+\mu\theta)(2+\mu\theta)}$ |
| $c_8$ | $-\frac{16}{256(1+\mu\theta)(2+\mu\theta)^2}$ | $\frac{16}{256(1+\mu\theta)(2+\mu\theta)^2}$ | $\frac{16}{256(1+\mu\theta)(2+\mu\theta)^2}$ | $\frac{32}{256(1+\mu\theta)(2+\mu\theta)^2}$ |
| $c_9$ | $\frac{2\mu\theta(4+\mu\theta)^2}{256(1+\mu\theta)^2(2+\mu\theta)^2(3+\mu\theta)}$ | $\frac{\mu\theta(32+40\mu\theta+10\mu^2\theta^2)}{256(1+\mu\theta)^2(2+\mu\theta)^2(3+\mu\theta)}$ | $\frac{2\mu^2\theta^2(4+\mu\theta)}{256(1+\mu\theta)^2(2+\mu\theta)^2(3+\mu\theta)}$ | $\frac{2\mu^2\theta^2(4+\mu\theta)}{256(1+\mu\theta)^2(2+\mu\theta)^2(3+\mu\theta)}$ |

|  | xyxz | yxxz | xyzw |
| --- | --- | --- | --- |
| $c_0$ | $\frac{1}{256}$ | $\frac{1}{256}$ | $\frac{1}{256}$ |
| $c_1$ | $-\frac{1}{256(1+\mu\theta)}$ | $-\frac{1}{256(1+\mu\theta)}$ | $-\frac{1}{256(1+\mu\theta)}$ |
| $c_2$ | $-\frac{2}{256(1+\mu\theta)}$ | $\frac{2}{256(1+\mu\theta)}$ | $-\frac{2}{256(1+\mu\theta)}$ |
| $c_3$ | $\frac{4}{256(1+\mu\theta)(2+\mu\theta)}$ | $-\frac{4}{256(1+\mu\theta)(2+\mu\theta)}$ | $\frac{4}{256(1+\mu\theta)(2+\mu\theta)}$ |
| $c_4$ | $\frac{1}{256(1+\mu\theta)}$ | $-\frac{3}{256(1+\mu\theta)}$ | $-\frac{3}{256(1+\mu\theta)}$ |
| $c_5$ | $-\frac{4}{256(1+\mu\theta)(2+\mu\theta)}$ | $\frac{4}{256(1+\mu\theta)(2+\mu\theta)}$ | $\frac{4}{256(1+\mu\theta)(2+\mu\theta)}$ |
| $c_6$ | $\frac{1}{256(1+\mu\theta)^2}$ | $\frac{1}{256(1+\mu\theta)^2}$ | $\frac{1}{256(1+\mu\theta)^2}$ |
| $c_7$ | 0 | 0 | $\frac{8}{256(1+\mu\theta)(2+\mu\theta)}$ |
| $c_8$ | 0 | 0 | $-\frac{16}{256(1+\mu\theta)(2+\mu\theta)^2}$ |
| $c_9$ | $-\frac{\mu^2\theta^2(4+2\mu\theta)}{256(1+\mu\theta)^2(2+\mu\theta)^2(3+\mu\theta)}$ | $-\frac{\mu^2\theta^2(4+2\mu\theta)}{256(1+\mu\theta)^2(2+\mu\theta)^2(3+\mu\theta)}$ | $\frac{2\mu^3\theta^3}{256(1+\mu\theta)^2(2+\mu\theta)^2(3+\mu\theta)}$ |

## Appendix B.

Below are the expressions for the variances of the parameters 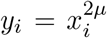,*i* = 1, 2, 3, for both the symmetric and the asymmetric species trees. In all of the expressions below, 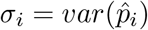 and 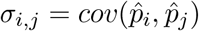, where *p*_*i*_ refers to the relevant entry in **p**_**s**_ or **p**_**a**_ for the symmetric or asymmetric species trees, respectively.

*B*.*1 Symmetric Case*

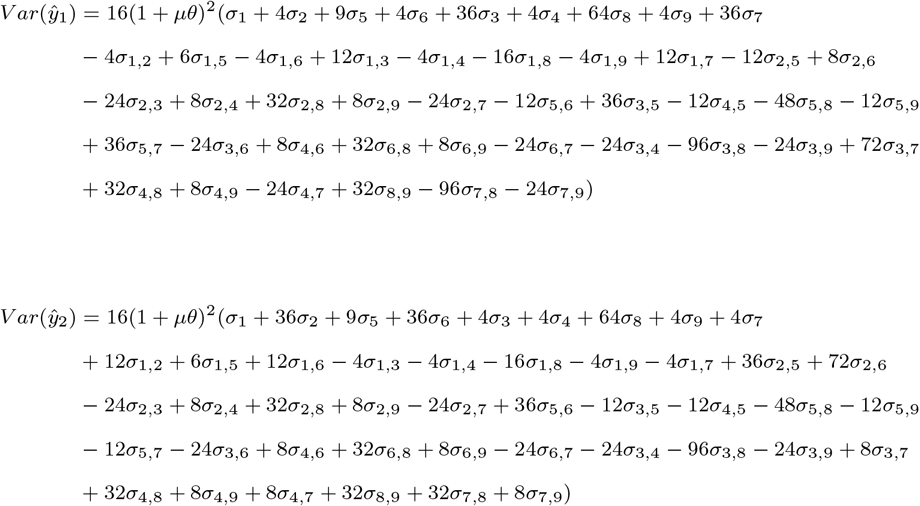

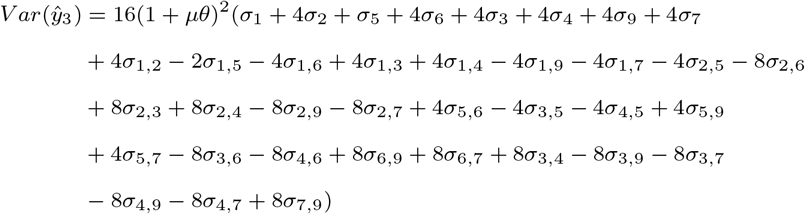

*B*.*2 Asymmetric Case*

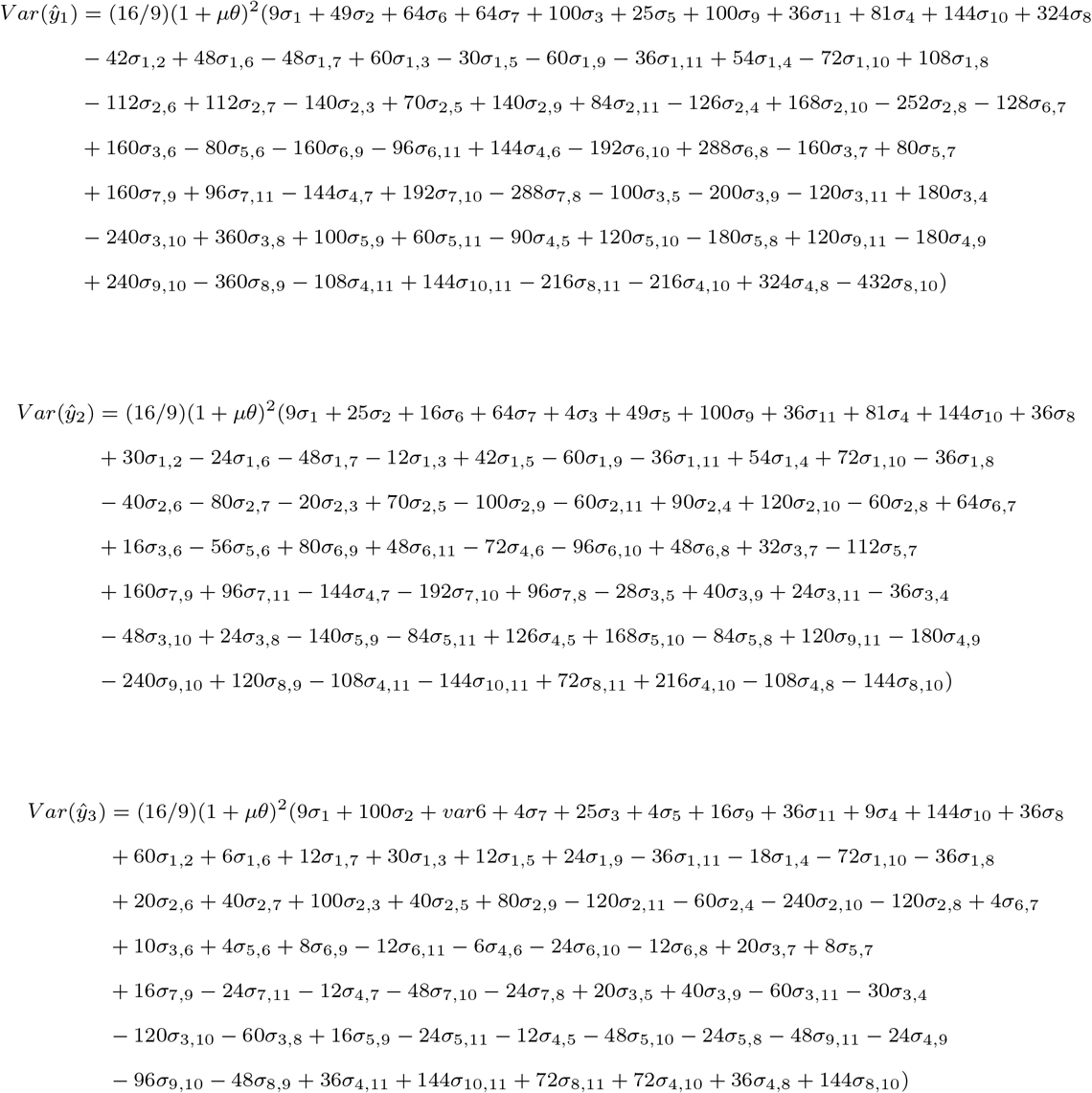

## Appendix C. Results for Three Taxa

In this section, we provide results analogous to those presented above for the case of three taxa. Our formulation draws on the work of Zhu and Yang (2021), who provided expressions for the site pattern probabilities of a three-taxon species tree under the JC69 model [10] in terms of the branch lengths and effective population size parameters. In their formulation, each branch was assigned a separate effective population size parameter. As we did in the case of four taxa, we assume that the effective population size is constant throughout the tree, and thus there is only a single effective population size parameter, *θ*.

We assume that the three-taxon species tree is of the form (a:*τ*_1_, b:*τ*_1_):*τ*_2_ *− τ*_1_,c:*τ*_2_);. We note that this notation matches what we used in the four-taxon case presented above, but differs from that used by Zhu and Yang (2021). There are 4 unique site pattern probabilities for three taxa, which we denote

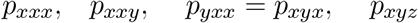

We can then solve Equation (18) in Zhu and Yang (2021) to get the following expressions:

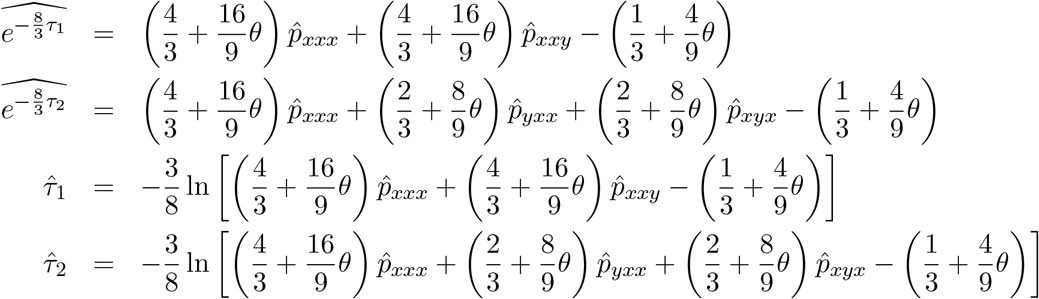

As in the case of four taxa, variances for the first two expressions can be derived and used to construct confidence intervals for *τ*_1_ and *τ*_2_. Following the same process as in the four-taxon case, we can derive the following expressions for the variances:

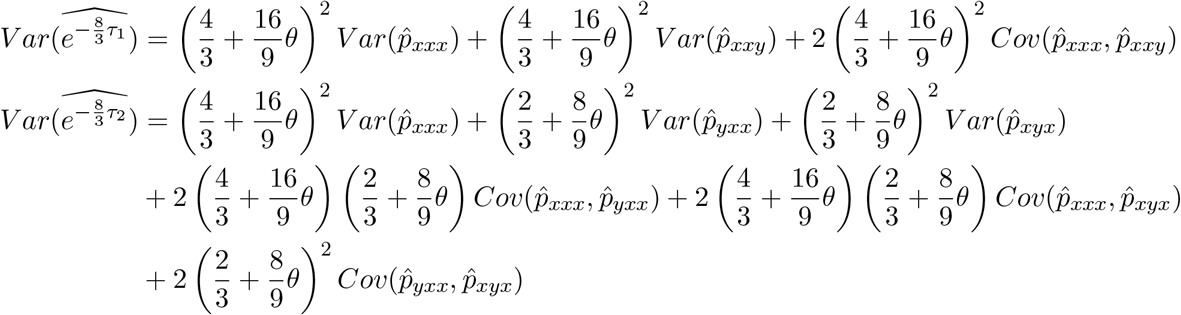

With the estimators of *τ*_1_, *τ*_2_, and their variances in hand, we can repeat the simulation studies undertaken in the four-taxon case. Specifically, we run simulations corresponding to Settings 1-4 in Section 4. We also considered the application to species delimitation described in Section 5. All results are presented in the Supplemental Material, and we include an R Markdown file with all of the code in the material available for download from GitHub.

## Notes

https://github.com/lkubatko/MOMSpeciationTimes

### Competing Interest Statement

The authors have declared no competing interest.

### Summary of Updates

Addition of an empirical example; minor re-wording to improve clarity and small notational changes.

